# The industrial anaerobe *Clostridium acetobutylicum* uses polyketides to regulate butanol production and differentiation

**DOI:** 10.1101/139873

**Authors:** Nicolaus A. Herman, Seong-Jong Kim, Jeffrey S. Li, Wenlong Cai, Hiroyuki Koshino, Wenjun Zhang

## Abstract

Polyketides are an important class of bioactive small molecules valued not only for their diverse therapeutic applications, but also for their role in controlling interesting biological phenotypes in their producing organisms. While numerous examples of polyketides derived from aerobic organisms exist, only a single family of polyketides has been identified from anaerobic organisms. Here we uncover a novel family of polyketides native to the anaerobic bacterium *Clostridium acetobutylicum*, an organism well-known for its historical use as an industrial producer of the organic solvents acetone, butanol, and ethanol. Through mutational analysis and chemical complementation assays, we demonstrate that these polyketides influence butanol production and act as chemical triggers of sporulation and granulose accumulation in this strain. This study represents a significant addition to the body of work demonstrating the existence and importance of polyketides in anaerobes, and showcases a novel strategy of manipulating the secondary metabolism of an organism to improve traits significant for industrial applications.

## INTRODUCTION

Polyketides are one of the most important classes of natural products given their wide range of applications in medicine and agriculture^1,2^. Encompassing several different chemical classes such as macrolides, polyenes, and aromatics, polyketides are employed for clinical use as antibiotics, anti-cancer agents, immunosuppressants, and even cholesterol-lowering drugs^3^. In addition to their role as therapeutic agents, many of these compounds are used by their producing organisms to access information about both the intracellular physiological status and extracellular environment, and control complex cellular processes such as morphological differentiation, virulence, stress response, and additional secondary metabolite production^4^. As virtually all known polyketides are derived from aerobic organisms (such as bacteria, fungi, and plants), there has been a long-standing assumption that anaerobic organisms are unable to produce these compounds^5^. However, recent genomic analysis has challenged this view by revealing that polyketide biosynthetic genes are widespread among anaerobic bacteria, and in particular, members from the genus *Clostridium*^6^. This diverse genus, comprised of anaerobic endospore-forming Gram-positive firmicutes, includes several notorious human pathogens as well as non-pathogenic species useful for industrial biotechnology^7^. To date, only one family of polyketides, the clostrubins, has been identified from *Clostridium*, representing the only known polyketides from the anaerobic world^8^. Clostrubins are pentacyclic polyphenolic polyketides biosynthesized by type II polyketide synthases (PKSs) in *C. beijerinckii^8^* and *C. puniceum^9^*, and display potent antibiotic activity against various pathogenic bacteria. Additionally, these aromatic polyketides have been shown to enable the plant pathogen *C. puniceum* to survive an oxygen-rich plant environment. The discovery of clostrubins provided the first experimental evidence that anaerobes are capable of producing bioactive polyketides, and motivated further studies on anaerobes to reveal additional polyketide metabolites with novel structures and interesting biological functions.

Here we have focused our attention on a type I modular PKS encoded on the genome of *C. acetobutylicum*, an organism historically used for industrial-scale production of the organic solvents acetone, *n*-butanol, and ethanol (ABE) through a process known as ABE fermentation^10^. Batch ABE fermentation by *C. acetobutylicum* ATCC 824 (the model ABE-producer) is characterized by two distinctive phases, an acid production phase (acidogenesis) and a solvent production phase (solventogenesis)^11^. During exponential growth, short-chain fatty acids (acetate and butyrate) are produced and accumulate in the media, causing a drop in the culture pH. As the culture approaches stationary phase, the previously formed acids are re-assimilated, the culture pH rises, and solvent production is initiated. The metabolic switch from acidogenesis to solventogenesis coincides with the initiation of the complex development program of sporulation^12^. As part of these processes, a starch-like carbohydrate known as granulose is produced and accumulates in swollen, phase-bright clostridial forms, within which endospores develop^13^. Further morphological development yields free spores, heat- and chemical-resistant cell types that do not contribute to solvent production.

Although ABE fermentation of *C. acetobutylicum* has been extensively studied, chemical signals responsible for triggering solventogenesis and/or sporulation have not been elucidated, and no secondary metabolite that plays a role in regulating these processes has been reported from this organism. Motivated by a recent transcriptomic analysis of *C. acetobutylicum*^14^ which showed that the expression of a type I modular PKS gene was significantly upregulated during early stationary phase (approximately 40-fold higher transcription level compared to mid-exponential phase), we postulated that the corresponding polyketide product might regulate one or more of the key fermentation phenotypes associated with solventogenesis and sporulation. Here we report the discovery of a family of polyketide metabolites using comparative, untargeted metabolomics followed by large-scale compound purification and molecular structure elucidation using NMR spectroscopy. Using a reverse genetics approach and chemical complementation, we further show that these polyketides influence butanol production, and are important in stimulating sporulation and granulose accumulation in *C. acetobutylicum*.

## RESULTS

### *In silico* analysis of the *pks* locus and inactivation of the *pks* gene in *C. acetobutylicum* ATCC 834

All sequenced *C. acetobutylicum* strains possess one PKS gene (*ca_c3355* in *C. acetobutylicum* ATCC 834) that encodes a type I single-module PKS with a predicted catalytic domain organization of ketosynthase, acyltransferase, dehydratase, ketoreductase, and acyl carrier protein (KS-AT-DH-KR-ACP) (**Supplementary Fig. 1a**). Homologous PKSs with the same domain structure could be identified, but the genes for these homologous PKSs are typically part of much larger biosynthetic gene clusters that encode additional PKSs and/or modification enzymes. *ca_c3355* appears to be an orphan gene with no other biosynthetic enzyme encoded either upstream or downstream, although several transcriptional regulators and efflux pumps are encoded in the vicinity (**Supplementary Fig. 1b**). While we were able to predict that malonyl-CoA was the likely substrate recognized by the AT domain (**Supplementary Fig. 1a**), it was difficult to predict the identity of polyketides which might be synthesized by this PKS, or if any chemical modifications of the nascent polyketide intermediate would be made by proteins encoded elsewhere on the genome. To determine the identity of any PKS-associated metabolites and probe the broader function of the *pks* locus, we performed a targeted in-frame deletion of the *pks* gene (*ca_c3355*) using an allelic exchange method previously developed for *C. acetobutylicum*^15^. The resulting mutant (Δ*pks*) was confirmed by PCR analysis (**Supplementary Fig. 2**).

### Identification of three polyketide metabolites through untargeted, comparative metabolomics

Quadruplicate batch fermentations with wild-type *C. acetobutylicum* and the mutant Δ*pks* were performed and harvested at early stationary phase. Organic extracts from combined supernatants and cell pellets were obtained and analyzed via liquid chromatography-high resolution mass spectroscopy (LC-HRMS). Following untargeted metabolomic comparisons of the two strains using XCMS^16^, three major species with molecular formulas C_14_H_22_O_3_ (**1**, calculated for C_14_H_21_O_3_^−^: 237.1496; found: 237.1496), C_20_H_32_O_7_ (**2**, calculated for C_20_H_31_O_7_^−^: 383.2075; found: 383.2080), and C_26_H_42_O_12_ (**3**, calculated for C_26_H_41_O_12_^−^: 545.2604; found: 545.2600) were found to be present in wild-type culture extracts and completely absent in extracts of Δ*pks* (**Fig. 1a** and **b; Supplementary Figs. 3-5**). A majority of **1**-**3** was found in the culture medium rather than cell pellets, suggesting that they were secreted into the culture medium upon production. **1**-**3** were UV active and featured identical UV absorption spectra (**Supplementary Figs. 3-5**), indicating that they share the same chromophore and likely the same biosynthetic origin. Furthermore, when the *pks* gene was introduced back into the Δ*pks* mutant, the genetic complementation strain was found to have resumed the production of **1**-**3**, demonstrating a direct relationship between the *pks* gene and the production of **1**-**3** (**Fig. 1b**).

**Figure 1.**
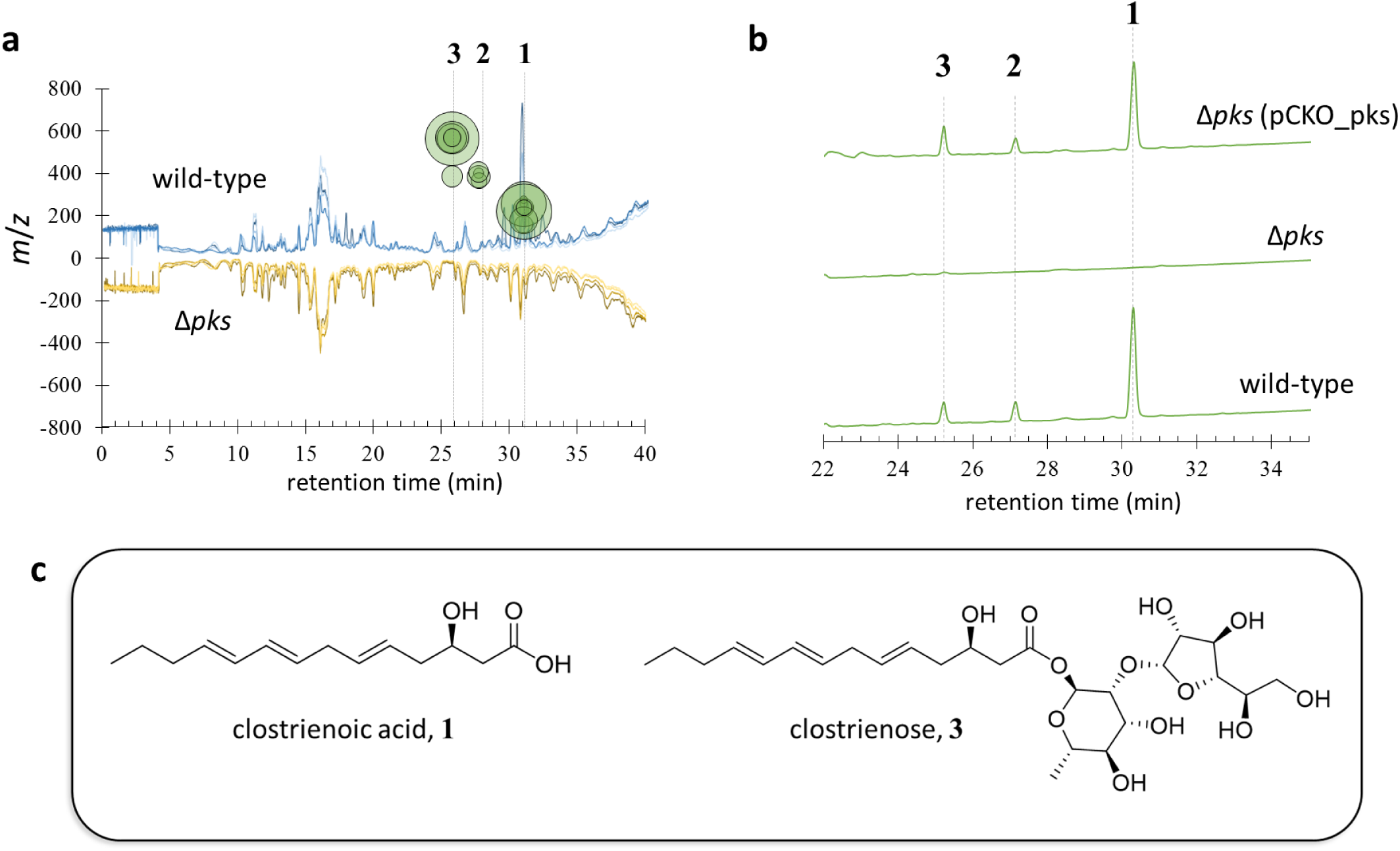
Identification of polyketides from wild-type extracts. **(a)** Results of XCMS analysis of wild-type *C. acetobutylicum* and Δ*pks* extracts taken from early stationary phase fermentation cultures. Blue (upper) chromatograms depict MS traces of quadruplicate wild-type extracts, while orange/brown (lower) chromatograms depict MS traces of quadruplicate Δ*pks* extracts. Green circles represent MS peaks unique to wild-type extracts, with corresponding *m*/*z* values indicated by the y-axis. No significant MS peaks were identified which were unique to Δ*pks*. The three peaks unique to wild-type are identified as **1**, **2**, and **3**. (**b**) UV-Vis traces (240 nm) of extracts from wild-type, Δ*pks*, and Δ*pks* genetic complementation strain [Δ*pks* (pCKO_pks)]. (**c**) Elucidated structures of **1** and **3** based on NMR characterization.

### Isolation and structure elucidation of polyketides

To isolate compounds **1**-**3**, we prepared 34 L of wild-type *C. acetobutylicum* culture broth. The culture was extracted with ethyl acetate and chromatographed on a silica gel column, followed by further purification via multiple rounds of HPLC using reverse-phase C18 columns. These purification steps yielded pure compound **1** (1.1 mg) and **3** (0.9 mg). NMR spectra, including ^1^H, ^13^C, dqf-COSY, TOCSY, HSQC, and HMBC spectra, were obtained for compound **1** (**Supplementary Table 1; Supplementary Fig. 6**). The proton signals at *δ*5.43, 5.56 and 6.00 indicated a considerable overlap in the aliphatic double bond region, and their configuration (all *E* stereochemistry) was further resolved using the high resolution HSQC spectrum without ^13^C decoupling. These assignments of geometric stereochemistry were also supported from ^13^C chemical shift values of allylic methylenes^17^. The carbon signals at *δ*172.97 and 67.25, together with related HMBC and TOCSY correlations, indicated the presence of a carboxylic acid and a secondary alcohol. The absolute stereochemistry of **1** was determined to be in the *R* configuration by measuring the specific rotation of its fully reduced product, 3-hydroxytetradecanoic acid (3-HTA). In particular, **1** was reduced to 3-HTA through hydrogenation over a palladium catalyst, and the specific rotation of the product 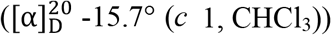 was consistent with the reported value of optically pure 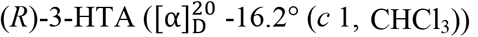^18^. From these data, we elucidated the molecular structure of **1** to be a modified tetradecenoic acid, which we termed clostrienoic acid (**Fig. 1c**). The predicted molecular formulas, MS/MS analysis, and identical UV absorption spectra suggested that **2** and **3** were related to **1**, likely containing additional monosaccharide and disaccharide moieties, respectively (**Supplementary Figs. 3-5**). Further 1D and 2D NMR spectroscopic analysis revealed that a disaccharide, α-D-galactofuranosyl(1->2)-α-L-rhamnopyranoside, is connected to the backbone of clostrienoic acid through an ester linkage in **3**, which we termed clostrienose (**Fig. 1c; Supplementary Table 2; Supplementary Fig. 7**). It is notable that the same disaccharide sequence has been found in several polysaccharides produced by microorganisms, and the relative configuration of each monosaccharide substituent in **3** was further confirmed by comparison to previously published NMR spectra^19–22^. An ester linkage between a sugar moiety and acid is rare, and only a few naturally occurring glycosyl esters have been previously reported, such as the *β*-D-glycosyl ester of 5-isoprenylindole-3-carboxylate isolated from *Streptomyces* sp. RM-5-8^23^, a microbial modification product of A58365A from *Streptomyces chromofuscus* NRRL 15098^24^, and a glycosyl ester of 3,4-seco-triterpene^25^. The minor compound **2** was indicated to contain one monosaccharide substituent (α-L-rhamnopyranoside) based on HRMS/MS analysis, but the titer was too low to be further confirmed by NMR spectroscopic analysis (**Supplementary Fig. 4**).

### *In vitro* biochemical analysis of PKS

Based on the structures of **1** and **3**, we hypothesized that the single-module PKS could use malonyl-CoA substrates and function iteratively to yield a heptaketide intermediate. Further modifications by putative auxiliary enzymes such as enoyl reductase, isomerase/desaturase, and thioesterase would then form the clostrienoic core (**1**). To confirm that *ca_c3355* encodes a functional iterative type I PKS, the 203 kDa megasynthase was overexpressed and purified from *E. coli* strain BAP1^26^. After incubating the purified PKS with ^14^C-labeled malonyl-CoA (generated *in situ* from 2-^14^C-malonic acid and CoA by malonyl-CoA synthase [MatB]^27^), the ^14^C-labeled covalent acyl-*S*-thiolation intermediate was detected by SDS-PAGE autoradiography (**Supplementary Fig. 8**). Further *in vitro* reconstitution of the activity of PKS with malonyl-CoA showed a dominant product, the triketide lactone **4**, which is a typical shunt product after spontaneous cyclic off-loading of the unreduced triketide. Addition of NADPH to this reaction enabled function of the KR domain (and the subsequent DH domain), leading to the production of the known shunt products tetraketide pyrone **5** and pentaketide pyrone **6** (**Supplementary Figs. 8** and **9**). Similar derailment in the normal programmed steps to yield the same shunt products has been observed in other iterative type I PKSs such as LovB^28^ and ApdA^29^. These results demonstrated that *ca_c3355* encodes a highly reducing type I PKS that functions iteratively, the first of its kind identified from an anaerobic organism. Furthermore, these results indicated that additional biosynthetic enzymes are needed for generating **1** and **3**, and their encoding genes are located elsewhere on the genome of *C. acetobutylicum*.

### Impact of the *pks* gene on ABE fermentation

To determine whether polyketide production influenced ABE fermentation, we compared the batch fermentation performance of wild-type *C. acetobutylicum* and Δ*pks* (**Fig. 2a-c; Supplementary Fig. 10**). While both strains displayed similar growth curves and the expected acidogenic and solventogenic phases, Δ*pks* showed stronger butanol production with ~10% increases in both butanol titer and productivity relative to wild-type (**Fig. 2a** and **b**). Perhaps related to the difference in butanol production, the butyrate concentration profiles also differed for Δ*pks* and wild-type; in the Δ*pks* culture, butyrate was produced more rapidly and reached a higher concentration during acidogenesis (0-20 hours), and was re-assimilated earlier during the transition to solventogenesis (20-23 hours) (**Fig. 2c**). This was also reflected in the pH profiles of the two strains, with the fall and rise of the culture pH (corresponding to the changes in metabolism) occurring earlier for Δ*pks* (**Fig. 2a**). Although less apparent, small increases in acetone and ethanol production were also observed for Δ*pks* relative to wild-type (**Fig. 2b**). These results established a clear link between the *pks* gene and ABE fermentation, and the production of polyketides in wild-type seemed to negatively affect the metabolic switch between the acidogenic and solventogenic phases.

**Figure 2.**
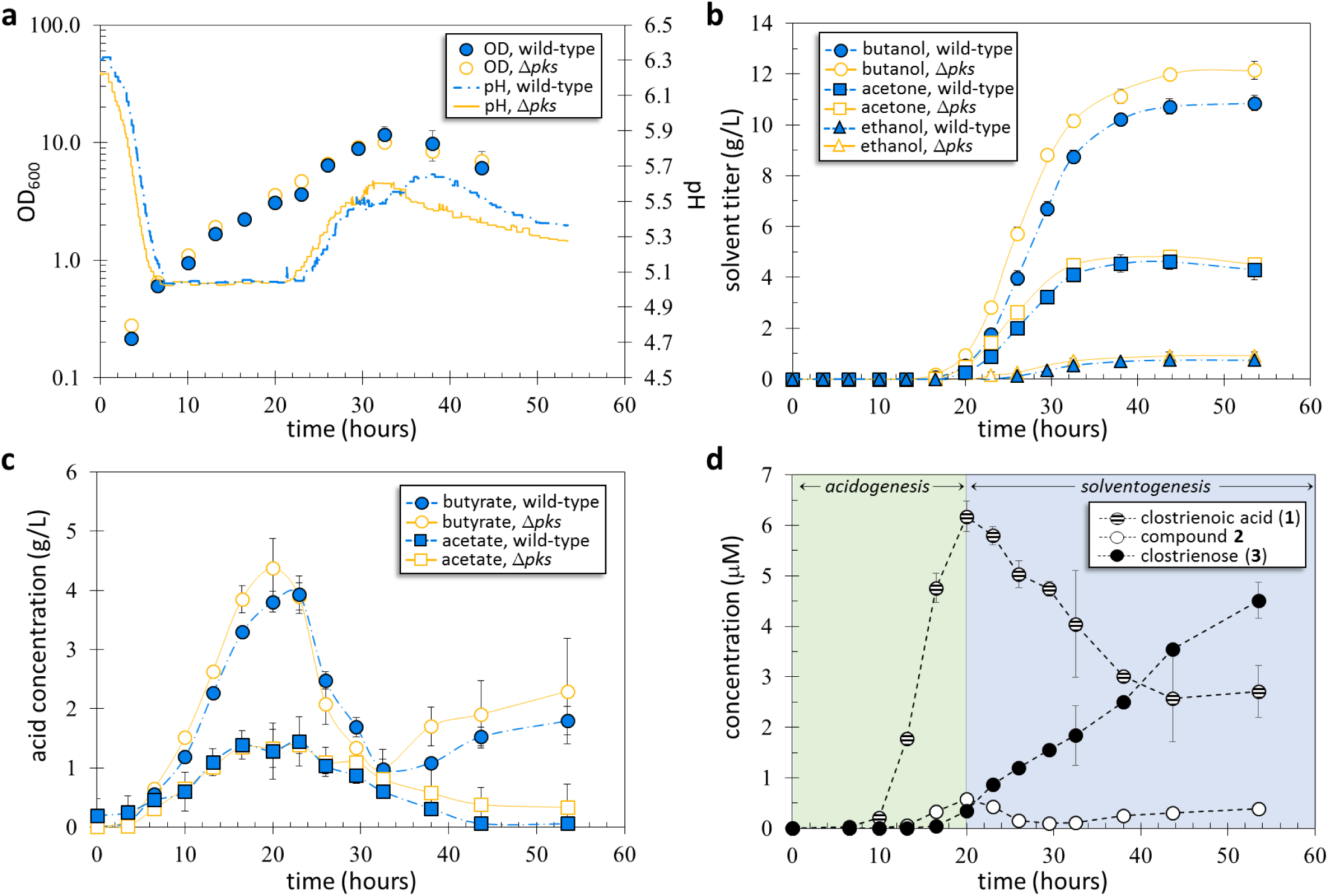
Time-course batch fermentation evaluation of wild-type *C. acetobutylicum* and Δ*pks*. **(a)** Culture density (measured as optical density at 600 nm, OD_600_) [left axis] and pH [right axis] from batch fermentations of wild-type and Δ*pks*. (**b**) Production of acetone, butanol, and ethanol from batch fermentations of wild-type and Δ*pks*. (**c**) Production of acetate and butyrate from batch fermentations of wild-type and Δ*pks*. (**d**) Production of compounds **1**-**3** from batch fermentation of wild-type *C. acetobutylicum* ATCC 824. As indicated, the green region represents the acid-production phase of the fermentation (acidogenesis), while the blue region represents the solvent-production phase (solventogenesis). Error bars represent the standard deviation of values from duplicate fermentations. Experiments were repeated at least three times independently.

To further probe the relationship between the production of the three polyketides and the ABE fermentation profile, we obtained production time-course profiles of **1**-**3** for wild-type *C. acetobutylicum* (**Fig. 2d**). Maximum levels of **1** (clostrienoic acid) and **2** were observed during the same period as maximum butyrate/acetate concentrations (20-23 hours), while the production of **3** (clostrienose) and butanol/acetone initiated at approximately the same time (~16 hours) and increased for the remainder of the fermentation. These results suggest a direct, although currently unclear, link between polyketide production and the ABE fermentation phases, with the production of **1** and **2** associating with acidogenesis, and the production of **3** associating with solventogenesis. We propose that **1** and **2** are biosynthetic intermediates of the end product **3**, and feeding studies with **1** and **2** showed that these compounds were readily converted to **3** in cultures of *C. acetobutylicum* Δ*pks*.

### Polyketides affect sporulation, granulose accumulation, and colony morphology

To better understand the broader biological impacts of polyketide production on *C. acetobutylicum* metabolism and physiology, we performed a transcriptome comparison of Δ*pks* and wild-type strains using RNA-Seq. Analysis of samples taken at early stationary phase revealed a total of 392 genes that were differentially expressed (expression fold change ≥ 2.0, p-value < 0.003) between Δ*pks* and wild-type, with 282 genes downregulated and 110 genes upregulated in Δ*pks*. STRING network analysis^30^ showed that the expression of genes related to four major cellular processes was downregulated in Δ*pks*, including sporulation (33 genes), carbohydrate transport (21 genes), carbohydrate metabolism (33 genes), and carbohydrate transfer (11 genes), and the expression of genes related to four other major cellular processes was upregulated in Δ*pks*, including sulfur metabolism (12 genes), cofactor biosynthesis (6 genes), stress response (8 genes), and amino acid transport (8 genes) (**Fig. 3; Supplementary Table 3**). It is notable that no significant difference in expression of the key solventogenic genes (including the crucial *sol* operon^31^) was observed, suggesting that the difference in butanol production between Δ*pks* and wild-type was not due to the direct transcriptional regulation of solventogenic genes by the polyketides. Consistent with the higher solvent production observed for Δ*pks*, the major pathways upregulated in Δ*pks* (particularly class I, III, and IV heat shock response machinery) have previously been associated with improved solvent tolerance in *C. acetobutylicum*^14,32,33^.

**Figure 3.**
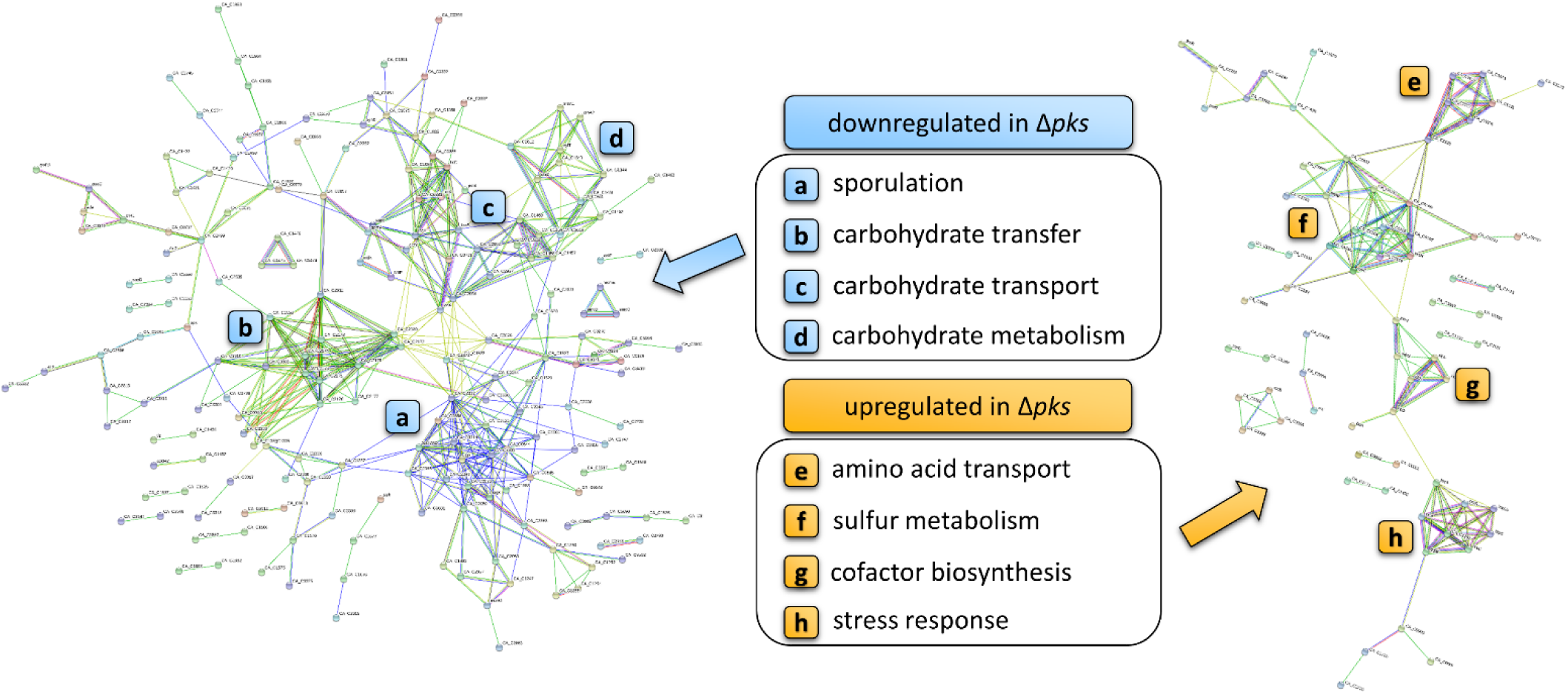
RNA-Seq comparison of wild-type *C. acetobutylicum* and Δ*pks*. STRING network analysis of genes predicted to be transcriptionally downregulated (left) and upregulated (right) by RNA-Seq in Δ*pks* fermentation culture relative to wild-type. Nodes represent differentially expressed genes, and lines represent predicted connections between genes including shared functional pathways of encoded proteins, chromosomal proximity, co-occurrence of genes in other organisms, co-expression of genes, and protein homology. For clarity, only genes with at least one predicted connection are shown. Four major concentrations of connected nodes were observed for each group (indicated by **a**-**h**), signifying the cellular pathways most affected by deletion of the *pks* gene. Detailed results from the RNA-Seq analysis are presented in **Supplementary Table 3**.

The sporulation genes downregulated in Δ*pks* include those encoding late stage sporulation proteins such as spore coat and germination proteins, and the sporulation-specific sigma factor K (σ^K^), one of the core regulators of sporulation in *C. acetobutylicum*^34^. Notably, transcription of the gene encoding the well-known master regulator of sporulation and solvent production, Spo0A^35^, was not significantly affected in Δ*pks*. To determine whether reduced sporulation was an observable property of Δ*pks* as suggested by the RNA-Seq analysis, sporulation assays in both liquid and solid media were performed for wild-type *C. acetobutylicum* and Δ*pks*. Indeed, a significant decrease in sporulation was observed for Δ*pks*, with spore formation decreasing by 3-4 orders of magnitude in liquid culture (**Fig. 4a**), and by ~2 orders of magnitude on solid media (*Supplementary Fig. 11*). Furthermore, the level of sporulation for Δ*pks* was partially restored when Δ*pks* culture broth was supplemented with clostrienose (**3**) (**Fig. 4a**). Since granulose biosynthesis and accumulation is related to the sporulation cycle, we then performed standard granulose accumulation assays using iodine staining. Following the trend observed for sporulation, a significant decrease in granulose accumulation was observed for Δ*pks*, and the addition of **3** to Δ*pks* culture completely restored granulose accumulation (even appearing to exceed levels observed for wild-type) (**Fig. 4b**). These assays revealed that the polyketides are important, although not essential, for triggering both sporulation and granulose accumulation in wild-type *C. acetobutylicum*. Interestingly, *pks* inactivation also affected colony morphology of *C. acetobutylicum*. While wild-type colonies were relatively flat in elevation and featured a distinctive “spore center” in the middle of the colony, Δ*pks* colonies were distinctively raised in elevation and featured a highly textured surface with no distinguishable spore center (**Fig. 4c**).

**Figure 4.**
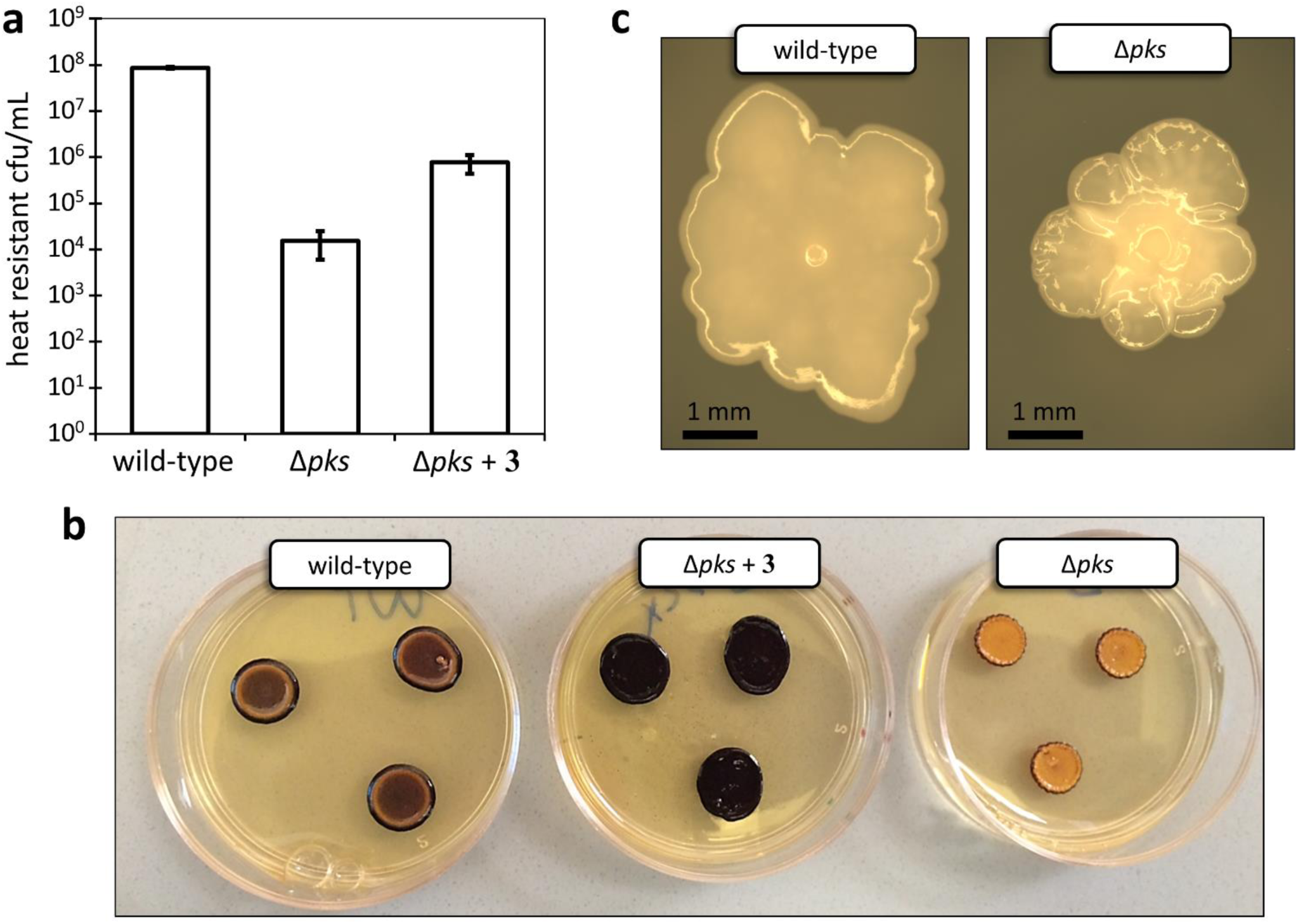
Phenotype comparison of wild-type *C. acetobutylicum* and Δ*pks*. **(a)** Sporulation assay results for liquid cultures of wild-type, Δ*pks*, and Δ*pks* supplemented with clostrienose (**3**). Values represent the number of heat resistant colony forming units per mL following incubation in liquid CBM-S medium for 5 days. Error bars represent the standard deviation of triplicate experiments. (**b**) Granulose accumulation assay results for wild-type, Δ*pks*, and Δ*pks* supplemented with clostrienose (**3**). Patches containing high levels of granulose are expected to stain dark brown/purple. (**c**) Dissecting microscope images of individual wild-type and Δ*pks* colonies.

## DISCUSSION

*Clostridium* is one of the largest bacterial genera, ranking second in size only to *Streptomyces*^36^. While members of *Streptomyces* are known to be prolific producers of secondary metabolites^37^, only a handful of secondary metabolites have been discovered from *Clostridium*. However, recent genomic analysis has indicated that secondary metabolite gene clusters can be found among diverse members of this genus, prompting efforts to identify and characterize these ‘cryptic’ secondary metabolites^6^. In this study, we identified a suite of polyketides (clostrienoic acid and clostrienose) from *C. acetobutylicum*, a well-studied solvent-producing anaerobe with no previously associated natural products.

Clostrienoic acid and clostrienose are biosynthesized by a predicted type I single-module PKS. Through *in vitro* reconstitution of the activity of purified PKS, we demonstrated that this megasynthase functions as a highly reducing iterative type I PKS, the first known of its kind among anaerobic organisms. We propose that the PKS functions iteratively to generate a heptaketide intermediate, which is then modified by tailoring enzymes encoded elsewhere on the genome to yield clostrienoic acid. Two subsequent glycosylation events are proposed to install first the rhamnopyranoside group, followed by the galactofuranoside group (generating clostrienose). The proposed sequential biosynthesis of the three polyketides is consistent with the production timing of **1**-**3** observed in batch fermentation of *C. acetobutylicum*. Although galactofuranose has previously been detected in cultures of *C. acetobutylicum*^38^ and the predicted biosynthetic genes for both sugar moieties are present on the *C. acetobutylicum* genome, the dedicated glycosyltransferases employed for the production of **3** are unclear. Due to the apparent non-clustered nature of genes encoding the enzymatic partners of PKS and the observation that hundreds of genes were upregulated along with *ca_c3355* at early stationary phase in *C. acetobutylicum*, extensive future work is needed to identify additional biosynthetic genes and elucidate the complete biosynthetic pathway of **3**.

Comparison of wild-type *C. acetobutylicum* and Δ*pks* along with chemical complementation of Δ*pks* strongly suggest that the identified novel polyketide, clostrienose (**3**), serves as a signaling molecule that regulates solventogenesis, granulose accumulation, and sporulation in *C. acetobutylicum*. Although cellular communication by signaling small molecules has been reported in a variety of Gram-negative and Gram-positive bacteria, with well-known examples such as *N*-acyl-homoserine lactones^39^, 4-quinolones^40^, γ-butyrolactones^41^, and autoinducer (AI)-2^42^, few signaling small molecules have been reported from *Clostridium*. In addition to clostrubin which helps the pathogenic *C. puniceum* access aerobic territory^9^, a putative AI-2-like system was reported to be involved in stimulating toxin production in *C. perfringens*^43^. Furthermore, gene clusters with homology to the well-studied *agr* quorum sensing system have been shown to be important for cellular processes such as toxin production^44–47^, biofilm formation^44^, and sporulation^47,48^ in several *Clostridium* species, although none of these studies identified the molecular structure of the presumed bioactive small molecules. This work provides solid evidence that morphological differentiation and solventogenesis can be regulated by secondary metabolites in *Clostridium*, and has revealed the molecular identity of the signal responsible for this behavior.

Solventogenic strains of *Clostridium* (such as *C. acetobutylicum*) were employed for industrial solvent production as early as 1916, and were eventually the source of 66% of US butanol production in 1945^10^. Although industrial operations of this process largely ceased during the 1950s due to the growth of the petrochemical industry, ABE fermentation has recently gained renewed interest given the wide range of agricultural feedstocks which can be converted to various commodity chemicals and potential biofuels using this process^49^. However, some drawbacks still exist that prevent the widespread use of ABE fermentation, such as low solvent titer/productivity due to solvent toxicity, and unfavorable cellular differentiation^50^. Rather than pursuing a traditional metabolic engineering strategy that focuses on the core metabolic pathway for solvent production, our work showcases an alternative approach by manipulating the secondary metabolism of the organism to improve traits significant for industrial ABE fermentation performance. In particular, given that both granulose biosynthesis and sporulation are undesirable traits for industrial fermentation (as granulose accumulation results in reduced solvent yields, and metabolically inactive spores do not contribute to solvent production)^50^, the reduced granulose accumulation and sporulation associated with Δ*pks* represent improved industrial traits. Furthermore, although none of the solvent producing genes were upregulated in Δ*pks* relative to wild-type, both butanol titer and productivity were increased in Δ*pks*. This may be explained by the decreased commitment of cells to sporulation in Δ*pks* (yielding a higher proportion of cells capable of solvent production), as well as the upregulation of cellular machinery related to butanol stress and adaptation as indicated by transcriptomic analysis.

In summary, we have discovered a family of novel polyketides that are biosynthesized by a highly reducing iterative type I PKS in *C. acetobutylicum* ATCC 824. In addition to the type II PKS-derived clostrubin, our work provides the second example of polyketide metabolites from a strictly anaerobic bacterium, and encourages continued efforts in exploring the uncharted terrain of secondary metabolites in the anaerobic world. We have further shown that the newly identified polyketides affect solvent production and are important for stimulating sporulation and granulose accumulation in *C. acetobutylicum*, adding to the extremely limited inventory of known signaling molecules used by *Clostridium* to control cellular physiology and metabolism. Furthermore, this work has yielded an engineered strain of *C. acetobutylicum* with improved traits for industrial ABE fermentation, demonstrating a novel strategy of manipulating secondary metabolism as a means of improving this important renewable bioprocess.

## Acknowledgements

We thank J. Liu (UC Berkeley) for performing the autoradiography labeling experiment, J. Pelton (UC Berkeley) for helping with NMR spectroscopic analysis, the University of California-Berkeley QB3 Functional Genomics Laboratory and Vincent J. Coates Genomic Sequencing Laboratory for performing the RNA-Seq library preparation and sequencing, and the Papoutsakis Lab (University of Delaware) for providing plasmids pKO_mazF and pAN1. This research was financially supported by the Energy Biosciences Institute, the Pew Scholars Program, the National Institutes of Health (DP2AT009148), and the Chan Zuckerberg Biohub Investigator Program.

## Author contributions

N.H. and W.Z. designed the experiments, analyzed the data, and wrote the manuscript. S.K. and W.C. performed purification and structure characterization of the polyketides. K.H. assisted with interpretation of NMR spectra of the polyketides. J.L. assisted with fermentation and phenotype characterization of *C. acetobutylicum* strains. N.H. performed all other experiments.

## Additional information

Supplementary information and chemical compound information are available in the online version of the paper. Reprints and permissions information is available online at http://www.nature.com/reprints/index.html. Correspondence and requests for materials should be addressed to W. Z.

## ONLINE METHODS

### Bacterial strains and media

*Clostridium acetobutylicum* ATCC 824 was cultured in an anaerobic chamber (Coy Laboratory Products) containing an atmosphere of 97% nitrogen and 3% hydrogen. 2xYTG medium^51^ contained 16 g/L tryptone, 10 g/L yeast extract, 5 g/L NaCl, and 10 g/L glucose (unless noted otherwise) with the pH adjusted to 5.2 for liquid media, and 5.8 for solid media (also containing 15 g/L agar). Clostridial Growth Medium (CGM)^52^ contained 30 g/L glucose (unless noted otherwise), 6.25 g/L yeast extract, 2.5 g/L ammonium sulfate, 1.25 g/L NaCl, 2.5 g/L asparagine, 0.95 g/L monobasic potassium phosphate, 0.95 g/L dibasic potassium phosphate, 0.5 g/L magnesium sulfate heptahydrate, 13 mg/L manganese sulfate heptahydrate, and 13 mg/L iron sulfate heptahydrate with the pH adjusted to 6.4. P2 medium^53^ contained 80 g/L glucose (unless noted otherwise), 1 g/L yeast extract, 2.2 g/L ammonium acetate, 0.5 g/L potassium phosphate monobasic, 0.5 g/L potassium sulfate dibasic, 0.2 g/L magnesium sulfate heptahydrate, 1 mg/L para-aminobenzoic acid, 1 mg/L thiamine hydrochloride, 10 μg/L biotin, 10 mg/L manganese sulfate heptahydrate, 10 mg/L ferrous sulfate heptahydrate, and 10 mg/L NaCl with the pH adjusted to 6.4. Clostridial Basal Medium (CBM)^54^ contained 10 g/L glucose, 0.5 g/L monobasic potassium phosphate, 0.5 g/L dibasic potassium phosphate, 4 g/L tryptone, 0.2 g/L magnesium sulfate heptahydrate, 10 mg/L manganese sulfate heptahydrate, 10 mg/L ferrous sulfate heptahydrate, 1 mg/L para-aminobenzoic acid, 1 mg/L thiamine hydrochloride, and 2 μg/L biotin with the pH adjusted to 6.9. For solid CGM plates, 15 g/L agar was added. CBM-S (used for liquid sporulation assays) was identical to CBM except 50 g/L glucose was used, and 5 g/L CaCO_3_ was added just prior to inoculation of cultures. *Escherichia coli* TOP10 (Thermo Fischer Scientific) was grown in Luria-Bertani (LB) medium at 37°C. For the appropriate *Clostridium* strains, culture media was supplemented with erythromycin (Ery; 40 μg/mL for solid media, 80 μg/mL for liquid media) and/or thiamphenicol (Th; 5 μg/mL for solid and liquid media). Kanamycin (Kan; 60 μg/mL) or chloramphenicol (Cm; 25 μg/mL) were added to *E. coli* culture media as indicated. *Clostridium* and *E. coli* strains were maintained as 20% v/v glycerol stocks stored at −80°C.

### Plasmid construction

Oligonucleotides were synthesized by Integrated DNA Technologies. **Supplementary Table 4** lists all relevant oligonucleotide sequences used. Phusion Polymerase (NEB) was used for all PCR amplifications. Genomic DNA isolation of *C. acetobutylicum* ATCC 824 was performed using a modified alkaline lysis method as previously reported^55^.

For constructing plasmid pKO_mazF_mod (which would later serve as a template for pKO_pks), primers pKON_Fo & pKON_Ro were used to PCR amplify a 5.0 kb region from the pKO_mazF template^15^. This step was necessary to remove a 677 bp region from the pKO_mazF backbone which we were unable to amplify via PCR. The 5.0 kb PCR product was gel purified, digested at the 5’ and 3’ ends using PshAI, ligated to form pKO_mazF_mod, and transformed into *E. coli* TOP10. Transformant clones were screened by purified plasmid test digestion, and Sanger sequencing was used to confirm the sequence of the final pKO_mazF_mod clone. For constructing plasmid pKO_pks (for deletion of the *pks* gene from *C. acetobutylicum*), a 3.8 kb region containing colE1, *repL, bgaR*, and *mazF* were PCR amplified from pKO_mazF_mod using primers pKO_F & pKO_R. Additionally, primers UHR_Fo & UHR_Ro and DHR_Fo & DHR_Ro were used to PCR amplify 1 kb regions representing the upstream and downstream homologous regions (UHR and DHR) flanking the *pks* gene (CA_C3355) from a *C. acetobutylicum* ATCC 824 genomic DNA template. The chloramphenicol/thiamphenicol resistance gene and constitutive promoter (P*ptb* from *C. acetobutylicum*) were PCR amplified from pKO_mazF_mod using primers CMR_F & CMR_R (1.1 kb region). These four PCR products were ligated via Gibson assembly and transformed into *E. coli* TOP10. Transformant clones were screened by purified plasmid test digestion, and Sanger sequencing was used to confirm the sequence of the final pKO_pks clone. For constructing plasmid pAN3^15^ (necessary for methylation of pKO_pks and pCKO_pks prior to transformation into *C. acetobutylicum)*, a 6.4 kb region from plasmid pAN1^56^ was amplified using primers pAN1_F & pAN1_R, and a 1.0 kb region containing the kanamycin resistance gene from vector pCOLA_Duet was amplified using primers KAN_Fo & KAN_Ro. The gel extracted PCR products were ligated via Gibson assembly and transformed in to *E. coli* TOP10. Transformant clones were screened by purified plasmid test digestion, and Sanger sequencing was used to confirm the sequence of the final pAN3 clone. For constructing plasmid pCKO_pks (for expression of the *pks* gene under the constitutive crotonase [*crt*] promoter from *C. acetobutylicum*), primers CPKS_Fo & CPKS_Ro were used to PCR amplify the 5.4 kb *C. acetobutylicum* ATCC 824 *pks* gene (CA_C3355) with appropriate overhangs for Gibson assembly. The pWIS_empty vector^57^ backbone (containing the constitutive *crt* promoter upstream of the multiple cloning site) was PCR amplified using primers pWIS_F & pWIS_R yielding a 5.0 kb product. The two gel purified PCR products were ligated via Gibson assembly and transformed into *E. coli* TOP10. Transformant clones were screened by purified plasmid test digestion, and Sanger sequencing was used to confirm the sequence of the final pCKO_pks clone.

For constructing plasmid pET24b_pks, the 5.4 kb *C. acetobutylicum* ATCC 824 *pks* gene (CA_C3355) was amplified from *C. acetobutylicum* genomic DNA using primers PKS_Fo and PKS_Ro. Doubly digested vector pET24b (EcoRI/XhoI digested) was ligated with doubly digested *pks* PCR product (EcoRI/XhoI digested), and the ligation product was transformed into *E. coli* TOP10. Transformant clones were screened by purified plasmid test digestion, and Sanger sequencing was used to confirm the sequence of the final pET24b_pks clone.

### Electro-transformation of *C. acetobutylicum*

Prior to transformation into *C. acetobutylicum*, vector pKO_pks was co-transformed with pAN3 into *E. coli* TOP10 via electroporation. This procedure permitted methylation of pKO_pks necessary for overcoming the native restriction-modification system active in *C. acetobutylicum*^56^. Plasmid purification of pKO_pks/pAN3 liquid culture was performed, and the resulting plasmid mixture was used for electroporations of *C. acetobutylicum*. Electroporations of *C. acetobutylicum* were performed as previously described^56^, with the exception that all transformation cultures were started from heat shocked (80°C for 10 minutes) single colonies.

### Deletion of *pks* gene (*ca_c3355*) and generation of genetic complementation strain

Targeted KO of the *pks* gene in *C. acetobutylicum* ATCC 824 was achieved using the previously published method^15^. Briefly, 5 μg of methylated pKO_pks/pAN3 plasmid mixture was transformed into *C. acetobutylicum* using the method referenced above. Following recovery in liquid 2xYTG medium for four hours, the cell pellets were collected by centrifugation, resuspended in 0.5 mL of fresh liquid 2xYTG, and 100 μL of the resuspended cell culture was plated on solid 2xYTG + 5 μg/mL Th + 40 mM β-lactose plates. Under these plating conditions, only cells which have undergone the desired double crossover homologous recombination event are expected to survive. Counterselection of the vector backbone is provided by the lactose-inducible promoter (P_*bgaL*_) which drives the toxin gene *mazF* on the pKO_pks vector backbone. Following this plating procedure, roughly 10 colonies were observed on the 2xYTG + 5 μg/mL Th + 40 mM β-lactose plates. Of these 10 colonies, four were twice restreaked and subjected to colony PCR verification. Four sets of primers were used as the basis of colony PCR verification, as detailed in **Supplementary Fig. 2**.

For generating the *pks* genetic complementation strain [Δ*pks* (pCKO_pks)], electroporation of Δ*pks C. acetobutylicum* with a pCKO_pks/pAN3 plasmid mixture was performed as described above. Following electroporation and recovery, cultures were plated on solid 2xYTG + 5 μg/mL Th + 40 μg/mL Ery media. After twice restreaking on solid 2xYTG + 5 μg/mL Th + 40 μg/mL Ery media, potential colonies harboring pCKO_pks were screened via colony PCR using primers pCKO_F & pCKO_R.

### LC-HRMS metabolomic analysis

For untargeted metabolomic comparisons of wild-type and Δ*pks C. acetobutylicum*, single colonies of each strain were heat shocked at 80°C for 10 min and used to inoculate 10 mL of liquid CGM (30 g/L glucose). Following overnight incubation (stagnant) until reaching OD_600_ ~ 1.0, these cultures were used to inoculate 10 mL of liquid CGM (80 g/L glucose) with a 10% inoculum for subculturing. After approximately five hours (OD_600_ ~ 1.0) of stagnant growth, the subcultures were used to inoculate quadruplicate flask fermentations (70 mL CGM, 80 g/L glucose + 5 μL Antifoam 204, 3% inoculum) agitated via magnetic stir bars. Calcium carbonate (6 g/L) was supplemented to the fermentation media for pH buffering. All culturing was performed at 37°C. 5 μg/mL Th was included in all cultures of Δ*pks* with the exception of the final fermentation culture, as certain antibiotics are known to perturb the ABE fermentation phases. Samples of fermentation broth (1 mL) from each replicate were taken during early stationary phase and extracted with 3 mL of 2:1 chloroform methanol. The mixtures were vortexed, separated via centrifugation (3,500 rpm, 10 min), and the bottom chloroform-rich layer was transferred to a glass vial. These organic extracts were dried with nitrogen gas, resuspended in 100 μL methanol, and 10 μL was injected onto an Agilent Technologies 6520 Accurate-Mass QTOF LC-MS instrument fitted with an Agilent Eclipse Plus C18 column (4.6 x 100 mm). A linear gradient of 2-98% CH_3_CN (vol/vol) over 40 min in H_2_O with 0.1% formic acid (vol/vol) at a flow rate of 0.5 mL/min was used. The metabolomic analysis platform XCMS^16^ (The Scripps Research Institute) was used to compare the metabolomes of wild-type and Δ*pks* strains based on the quadruplicate LC-HRMS data. MS peaks unique to Δ*pks* (**Fig. 1a**) were identified using the following parameters: p-value < 0.01, fold change > 10.0, peak intensity > 5000.

### Isolation, purification, and structure elucidation of compounds 1, 2 and 3

To yield sufficient amounts of compounds **1**, **2**, and **3** for complete structure elucidation, 34 L of wild-type *C. acetobutylicum* culture was generated from several rounds of fermentation using a New Brunswick Scientific Bioflow 115 Benchtop Fermenter. Overnight cultures (10 mL CGM, 30 g/L glucose, stagnant, 37°C) inoculated with heat shocked individual colonies of *C. acetobutylicum* were cultured until reaching OD_600_ ~ 1. A 10% inoculum was then used to start a subculture (50 mL CGM, 30 g/L glucose, stagnant, 37°C), and the subculture was incubated until reaching OD_600_ ~ 1. A 10% inoculum was used to start the final subculture (300 mL CGM, 80 g/L glucose, stagnant, 37°C), and the final subculture was incubated until reaching OD_600_ ~ 1. The 300 mL culture was then aseptically transferred to the Bioflow 115 fermenter pre-loaded with 10.0 L of nitrogen-sparged CGM (80 g/L glucose, 1 mL Antifoam 204, 37°C). The fermentation was then allowed to proceed for 72 hours. The temperature was maintained at 37°C, agitation was provided by stirring at 200 rpm, and the pH was maintained above 5.0 via automatic addition of 3 M NH_4_OH.

After combining the pooled fermentation culture (34 L) and removing the cell pellets via centrifugation, the cell-free culture supernatant was extracted using two volumes of ethyl acetate. Following isolation of the organic extract, the solvent was removed by rotary evaporation and the residue was redissolved in dichloromethane. The dark yellow oily residue (10.8 g) was subjected to silica gel column chromatography (60 Å, 220-440 mesh), and the compound-rich fractions were eluted with an ethyl acetate/hexane gradient system. Each of the compound-rich fractions were combined and concentrated to dryness (3.6 g of compound **1**, 1.1 g of compound **2**, and 2.4 g of compound **3**). Further purification of compounds **1**, **2** and **3** was performed individually. Each yellow residue of compounds **1**, **2** and **3** was subjected to reverse-phase high-performance liquid chromatography (RP-HPLC, Agilent 1260 HPLC with DAD) on a semi-preparative C18 column (10 μm, 250 x 22 mm, 300 Å, Vydac) with a linear gradient of 2-98% CH_3_CN (vol/vol) over 1 hour in H_2_O with 0.1% formic acid at a flow rate of 5 mL/min. Product-rich fractions containing compound **1**, **2** and **3**, respectively, were collected manually and concentrated under vacuum. The residue was redissolved in methanol (2-4 mL), and was again purified by RP-HPLC (Agilent 1260 HPLC with DAD) using a C18 column (5 μm, 250 x 10 mm, 300 Å, Vydac) with a linear gradient of 25-80% CH_3_CN (vol/vol) over 30 min in H_2_O with 0.1% formic acid (vol/vol) at a flow rate of 2 mL/min. Fractions containing compound **1**, **2** and **3**, respectively, were collected manually and concentrated under vacuum. The residues of compound 1, 2 and 3 were redissolved in methanol (0.5-2 mL) for further purification. Purification was again performed by RP-HPLC using a C18 column (5 μm, 250 x 4.6 mm, 100 Å, Inertsil ODS-4 column) with an isocratic system of 22% (compound **3**), 25% (compound **2**), and 30% (compound **1**) CH_3_CN (vol/vol) over 30 min in H_2_O with 0.1% formic acid (vol/vol) at a flow rate of 1 mL/min. Fractions containing compound **1**, **2** and **3**, respectively, were collected manually and concentrated under vacuum. The resulting purified compounds **3** were dried and analyzed by HRMS and NMR. The titer of **2** was too low for NMR analysis. The residue of compound **1** required further purification, and was redissolved in methanol (0.5 mL) again for the final purification. The final purification was performed by RP-HPLC using a C18 column (3.5 μm, 100 x 4.6 mm, Agilent Eclipse Plus) with an isocratic system of 26% CH_3_CN (vol/vol) over 30 min in H_2_O with 0.1% formic acid (vol/vol) at a flow rate of 0.5 mL/min. Fractions containing compound **1** were collected manually and concentrated under vacuum. The resulting purified compound **1** was dried and analyzed by HRMS and NMR. LC-HRMS analysis was performed on an Agilent Technologies 6520 Accurate-Mass QTOF LC-MS instrument fitted with an Agilent Eclipse Plus C18 column (4.6 x 100 mm). A linear gradient of 2-98% CH_3_CN (vol/vol) over 40 min in H_2_O with 0.1% formic acid (vol/vol) at a flow rate of 0.5 mL/min was used. NMR spectra were recorded on a Bruker Biospin 900 MHz spectrometer with a cryoprobe in d4-methanol or d6-DMSO (Cambridge Isotope Laboratories).

To determine the absolute stereochemistry of the secondary alcohol in **1**, an additional 15 mg of **1** was purified and dissolved in 10 ml of absolute EtOH, and 0.75 mg of 10% Pd/C was added. The flask was purged with hydrogen, and the reaction was stirred at room temperature for 3 hours under positive pressure of hydrogen. Reaction progress was monitored by TLC. The reaction was quenched by filtration, and the solvent was dried under vacuum to yield 3-HTA as a white powder (reaction yield: 75%). The resulting 3-HTA product was analyzed by LC-HRMS, ^1^H NMR, ^13^C NMR, and the specific rotation was measured. ESI-HRMS (m/z) = 243.1933 ([M-H]^−^) (calc. 243.1960). ^1^H NMR (900 MHz, CDCl_3_) δ 4.01 (m, 1H), 2.56 (dd, J = 16.6, 3.0 Hz, 1H), 2.46 (dd, J = 16.6, 9.1 Hz, 1H), 1.53 (m, 1H), 1.44 (m, 2H), 1.35 – 1.22 (m, 17H), 0.86 (t, J = 7.2 Hz, 3H). ^13^C NMR (226 MHz, CDCl_3_) δ 176.28, 68.22, 40.95, 36.76, 32.13, 29.86, 29.84, 29.79, 29.77, 29.69, 29.56, 25.65, 22.91, 14.35. 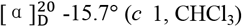. The specific rotation of **1** was also measured 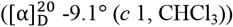.

### Bioreactor fermentations

To compare ABE fermentation profiles of wild-type and Δ*pks C. acetobutylicum*, bioreactor fermentations were carried out in DASGIP Bioreactors (4 x GPI 100 Vessels, DASGIP Bioblock System) with 500 mL working volumes. Overnight cultures (10 mL CGM, 30 g/L glucose, stagnant, 34°C) inoculated with heat shocked individual colonies of *C. acetobutylicum* were cultured until reaching OD_600_ ~ 1. A 10% inoculum was then used to start a subculture (30 mL P2 media, 80 g/L glucose, stagnant, 34°C), and the subculture was incubated until reaching OD_600_ ~ 1. The 30 mL subcultures were then aseptically transferred into individual DASGIP Bioreactors pre-loaded with 500 mL P2 medium (80 g/L glucose, 100 μL Antifoam 204, 34°C). The fermentations were allowed to proceed for 54 hours with periodic sampling for optical density measurements, fermentation product analysis, and quantification of compounds **1**, **2**, and **3**. The temperature was maintained at 34°C throughout the fermentation, agitation was provided by stirring at 200 rpm, and the pH was maintained above 5.0 via automatic addition of 3 M NH_4_OH. To maintain anaerobic conditions, oxygen-free nitrogen gas was sparged at a rate of 2 sL/h for the duration of the fermentation. For quantification of compounds **1**, **2**, and **3**, 1 mL of fermentation broth was mixed with 3 mL ethyl acetate, vortexed, separated via centrifugation (3,500 rpm, 10 min), and the upper organic layer isolated. The organic layer was dried by rotary evaporation, resuspended in 200 μL methanol, and 20 μL was injected onto an Agilent Technologies 6120 Quadrupole LC-MS (with DAD) instrument fitted with an Agilent Eclipse Plus C18 column (4.6 x 100 mm). A linear gradient of 2-98% CH_3_CN (vol/vol) over 40 minutes in H_2_O with 0.1% formic acid (vol/vol) at a flow rate of 0.5 mL/min was used. Compounds **1**, **2**, and **3** were identified by UV absorption (240 nm) as demonstrated in **Fig. 1b**, and were quantified by the integrated peak area (absorbance at 240 nm).

### Fermentation analytical procedures

Cell concentrations were monitored via absorbance using a spectrophotometer (optical density at 600nm, OD_600_). Glucose, acetone, butanol, ethanol, acetate, butyrate, and lactate concentrations were measured using a Shimadzu Prominence UFLC system with refractive index and diode array detection. Prior to analysis, samples of culture supernatant were filtered with 0.22 micron PVDF syringe filters. Compounds were separated with a Biorad Aminex HPX-87H column (300 mm x 7.8 mm) and detected by measurement of refractive index (glucose, butanol, ethanol, acetate, lactate) or UV absorbance (acetone, 265 nm; butyrate, 208 nm). The following operating conditions were used: mobile phase, 0.01 N sulfuric acid; flow rate, 0.7 mL/min; run time, 35 min; column temperature, 35°C.

### RNA isolation and RNA-Seq analysis

Samples (10 mL) of fermentation broth were taken in triplicate from bioreactor fermentations of wild-type and Δ*pks C. acetobutylicum* 26 hours post-inoculation. The samples were centrifuged (4,000 xg, 10 min, 4°C) and the pellets were resuspended and stored in RNAprotect Cell Reagent (Qiagen). Total RNA was extracted using an RNeasy Mini kit (Qiagen) according the manufacturer’s instructions. An on-column DNase treatment was performed using DNase I (RNase-free) (NEB). RNA quality control, library construction, and library sequencing were performed by the University of California-Berkeley QB3 Functional Genomics Laboratory and Vincent J. Coates Genomic Sequencing Laboratory. RNA quality and concentration was assessed using a nanochip on an Agilent 2100 Bioanalyzer. Bacterial 16S and 23S rRNA was removed using a RiboZero kit (Illumina). The resulting mRNA was converted to an RNA-Seq library using an mRNA-Seq library construction kit (Illumina). RNA library sequencing was performed on an Illumina HiSeq4000 with 50 bp single end reads. Sequencing reads (50 bp) were processed and mapped to the *C. acetobutylicum* ATCC 824 genome (NCBI accession NC_003030.1 [chromosome] and NC_001988.2 [megaplasmid]) using CLC Genomics Workbench 9.0 with default parameters. Reads that did not uniquely align to the genome were discarded. Differences in gene expression between wild-type and Δ*pks C. acetobutylicum* were calculated using the same software. Genes were considered differentially expressed with p-value < 0.003 and | normalized fold change | ≥ 2.0. Results of the RNA-Seq analysis are presented in **Supplementary Table 3**. STRING analysis^30^ was performed to determine putative protein-protein interactions between the differentially expressed genes revealed by RNA-Seq analysis.

### Sporulation assays, granulose accumulation assay, and light microscopy

Liquid sporulation assays were performed as previously described^58^ with minor modifications. Samples were taken from biological triplicate liquid cultures after five days of incubation (30 mL CBM-S, 37°C). The 20 μL samples were heat shocked (80°C, 10 min), dilutions (10^1^-10^6^) were spotted on 2xYTG plates, and colonies were enumerated after 30 hours of incubation (37°C) to calculate the number of heat resistant colony forming units (cfu/mL). For chemical complementation of Δpks in the liquid sporulation assay, purified clostrienose (final concentration 3.5 μM) was supplemented in CBM-S cultures of Δ*pks* C. *acetobutylicum* at the time of inoculation. Since compound **3** was added as a concentrated solution in methanol, the equivalent volume of methanol (60 μL) was added to all other liquid sporulation assay cultures (wild-type and non-complemented Δ*pks*) to control for this effect.

For solid sporulation assays, a previously described method^59^ was employed with some modifications. Heat shocked (80°C, 10 min) individual colonies were cultured in liquid media (10 mL CGM, 37°C) for 24 hours. Cultures were diluted by a factor of 10^6^ and plated on solid CBM. Sampling was conducted 1, 2, 3, 4, and 6 days following initial plating. For sampling, three individual colonies were combined and thoroughly resuspended in 60 μL of liquid CGM. 20 μL of the resuspended colony mixture was then heat shocked (80°C, 10 min), dilutions (10^1^-10^6^) were spotted on 2xYTG plates, and colonies were enumerated after 30 hours of incubation (37°C) to calculate the number of heat resistant colony forming units (cfu/colony). All samples were performed as biological triplicates.

Granulose accumulation assays were performed via iodine staining as previously described^48^ with some modifications. Mid-log phase (OD_600_ ~ 0.6) liquid cultures (P2, 37°C) were plated on solid 2xYTG medium with elevated glucose levels (50 g/L) to enable granulose production. The plates were incubated at 30°C for four days, at which point they were stained by exposure to a bed of iodine crystals for 10 minutes. The plates were then allowed to destain for 10 minutes prior to imaging. For chemical complementation of Δ*pks* in the granulose accumulation assay, purified clostrienose (final plate concentration 4.0 μM) was embedded in solid 2xYTG plates prior to plating of Δ*pks* culture. Since clostrienose was added to 2xYTG plates as a concentrated solution in methanol (mixed into the molten media prior to solidification), the equivalent volume of methanol (6 μL) was added to all other 2xYTG plates used in the granulose accumulation assays to control for this effect.

Individual colonies of *C. acetobutylicum* cultured on CBM plates for 48 hours (37°C) were viewed and imaged using a Leica MZ16 F dissecting microscope fitted with a Leica DFC300 FX camera.

### Purification and *in vitro* analysis of PKS protein

To purify the His6-tagged PKS protein for *in vitro* analysis, pET24b_pks was transformed into *E. coli* BAP1. A single colony was inoculated into 10 mL LB + 50 μg/mL kanamycin (Kan) for overnight growth at 37°C. 7 mL of overnight culture was used to inoculate 700 mL LB + 50 μg/mL Kan, and the culture was shaken at 240 rpm and 37°C until reaching OD_600_ of 0.5. After icing the culture for 10 minutes, isopropyl thio- β -D-galactoside (IPTG) was added to a final concentration of 0.1 mM to induce protein expression, and the culture was incubated at 16°C for 16 hours. The cells were harvested by centrifugation (6,000 rpm, 4°C, 20 minutes), resuspended in 25 mL lysis buffer (25 mM HEPES, pH 8.0, 0.5 M NaCl, 5 mM imidazole), and lysed by homogenization over ice. Cell debris was removed by centrifugation (15,000 rpm, 4°C, 60 minutes), and Ni-NTA agarose resin was added to the supernatant (2 mL/L culture). The mixture was nutated at 4°C for 1 hour, loaded onto a gravity flow column, and the PKS protein was eluted with increasing concentrations of imidazole in Buffer A (20 mM HEPES, pH 8.0, 1 mM DTT). Purified PKS protein was concentrated and buffer exchanged into Buffer A + 10% glycerol using a Vivaspin Centrifugal Concentrator. Aliquots of purified PKS protein were aliquoted and flash frozen in liquid nitrogen. The approximate protein yield was 5 mg/L (203 kDa).

A PKS loading assay with ^14^C-labeled substrate contained, in a total volume of 15 μL, 4 mM ATP, 2 mM MgCl_2_, 1 mM TCEP, 0.5 mM CoA, 2-^14^C-malonic acid [0.1 μCi], 10 μM MatB, 17 μM PKS, and 50 mM HEPES, pH 8.0. After 2 hours of incubation at 25 °C, samples were quenched by adding an equal volume of 1× SDS sample buffer. Following SDS-PAGE analysis with a 4-15% TGX gel (Criterion), the gel was dried for 2 hours at 50°C and exposed on a storage phosphor screen (20 x 25 cm; Molecular Dynamics) for 5 days. Radiolabeled protein was imaged on a Typhoon 9400 phosphorimager (Storage Phosphor mode, 50 μm resolution; Amersham Biosciences).

For *in vitro* product assays (50 μL) of the PKS protein, 8 μM of PKS was incubated with malonyl-CoA (2 mM) in phosphate buffer (100 mM, pH 7.0) at room temperature for 2 hours to generate compound 4. NADPH (2 mM) was added to the assay to generate compounds 5 and 6. Following incubation, the assay mixtures were extracted twice with 100 μL of 99% ethyl acetate/1% acetic acid (v/v). The organic extracts were dried and resuspended in 100 μL of methanol, and 10 μL was injected onto an Agilent Technologies 6120 Quadrupole LC-MS (with DAD) instrument fitted with an Agilent Eclipse Plus C18 column (4.6 x 100 mm). A linear gradient of 2-98% CH_3_CN (vol/vol) over 40 minutes in H_2_O with 0.1% formic acid (vol/vol) at a flow rate of 0.5 mL/min was used. LC-HRMS analysis of the assay extracts was performed on an Agilent Technologies 6520 Accurate-Mass QTOF LC-MS instrument fitted with an Agilent Eclipse Plus C18 column (4.6 x 100 mm) using the same solvent gradient and flow rate described above.

